# Advancing EEG Biosensing: Novel Scalable Production of Platinum-Silicon Microneedles

**DOI:** 10.1101/2025.03.29.646121

**Authors:** Momina Amir, Ruochen Ding, Nurul Izni Rusli, Nadalan Vercooren, Frederik Ceyssens, Nadezda Kuznetsova, Alexander Bertrand, Michael Kraft, Irene Taurino

## Abstract

This study presents the development and analysis of a silicon microneedle electrode array fabricated using a streamlined, two-step approach for enhanced bio-signal recording applications. Numerical simulations with COMSOL Multiphysics were employed to optimize the microneedle array design, focusing on mechanical stability and minimizing skin-electrode impedance. Structural analysis identified needle lengths between 500 μm and 700 μm as optimal for mechanical robustness, with critical load factors indicating a high resistance to buckling under applied forces. Optimal needle spacing was determined to prevent inter-needle interference and ensure effective skin penetration. Proof-of-concept electroencephalography testing confirmed that the microneedles achieved reliable bio-signal recording, comparable to traditional wet electrodes, while offering advantages such as reduced preparation time and improved user comfort. This work establishes a simple and cleanroom-compatible, scalable method for fabricating platinum-coated silicon microneedle arrays, advancing the development of wearable, user-friendly biopotential monitoring devices.

## Introduction

In recent years, dry electrodes have attracted widespread interest in biomedical applications due to their ability to interact with the body in a minimally invasive manner, reducing discomfort and the risk of infection. These electrodes provide a versatile platform for a wide range of medical diagnostics and therapeutic interventions. Among these technologies, microneedle (MN) array electrodes have emerged as a particularly promising option for applications such as transdermal drug delivery [1] and bio-signal monitoring, including electroencephalography (EEG), electrocardiography (ECG), and electromyography (EMG) [2]. Moreover, MN electrodes have been effectively utilized in tissue stimulation, aiding in regeneration and healing while complementing their functions in drug delivery and signal monitoring. This versatility highlights their potential to advance biomedical technologies and broaden their scope of applications in healthcare [3].

Despite their versatility, the performance of MN electrodes is often constrained by critical technical challenges, primarily related to impedance and skin penetration. The stratum corneum (SC), the outermost layer of the skin, serves as a protective barrier but also poses high impedance, hindering signal acquisition and drug delivery. MN electrodes overcome these obstacles effectively by bypassing the SC. Designed with sufficient stiffness to penetrate the outer skin layer, they provide direct access to underlying tissues, significantly reducing impedance and ensuring minimal signal distortion for more accurate signal acquisition [2], [4], [5].

The mechanical design of the MNs plays a pivotal role in their effectiveness, as the needles must possess sufficient strength to penetrate the SC without undergoing significant deformation or breakage. Recent studies have emphasized optimizing parameters such as needle length, pitch, and structural stability to achieve a balance between mechanical robustness and penetration efficiency [6], [7]. While materials such as polymers and metals have been explored for MN fabrication, silicon (Si) remains a preferred choice due to its high mechanical strength, precise manufacturability, and resistance to wear and fatigue. Additionally, its favorable Young’s modulus and compatibility with complementary metal-oxide semiconductor (CMOS) technologies make it especially suitable for advanced applications [8], [9], [10].

Various techniques for fabricating Si MNs have been developed, including laser machining, photolithography, and wet and dry etching [9], [10], [11], [12]. The choice of fabrication method significantly impacts the performance and scalability of Si MN arrays. In contrast to other methods, wet etching with potassium hydroxide (KOH) is often preferred due to its simplicity, scalability, and ability to produce precise anisotropic structures with minimal processing complexity [8][10]. This technique exploits the crystallographic anisotropy of Si to create MNs with sharp tips and well-defined geometries, ensuring effective skin penetration and reduced impedance. Unlike more complex methods, such as photolithographic patterning or deep reactive ion etching (DRIE), wet etching minimizes fabrication steps, reduces mechanical stress on the Si substrate, and is compatible with cleanroom environments, making it a practical and efficient solution for fabricating robust, scalable Si MN arrays.

This paper presents a streamlined two-step fabrication process for a Si MN electrode array that involves wet etching in KOH solution on a partially diced wafer. Structural analyses using COMSOL finite element modeling were conducted prior to fabrication to determine parameters that ensure mechanical robustness, including optimal needle height and spacing (pitch). The fabricated MN arrays were evaluated for their biosignal acquisition capabilities through proof-of-concept EEG recordings, demonstrating their potential as user-friendly alternatives to conventional wet electrodes.

## Simulations

To optimize the structural design of the Si MNs and determine the optimal dicing parameters, finite element analysis (FEA) was conducted using COMSOL Multiphysics. The simulation protocol followed by Loizidou et al. [13] was adapted for modeling a 3×3 Si MN array on a Si substrate. The skin was approximated as two cylindrical layers—the epidermis and dermis—with material and geometric properties listed in Table 1. A force of 5 N, based on prior studies [13][14], was applied to simulate skin penetration under realistic conditions. Although this two-layer model simplifies the inherent heterogeneity of human skin, the validation work by Loizidou et al. [13] has shown it to be sufficiently representative for comparing different microneedle lengths and pitches under consistent conditions.

**Table 1.** Material and geometric parameters.

| **Silicon** |  |
| --- | --- |
| Density | 2.39 g/cm<sup>3</sup> |
| Young's Modulus | 130 GPa |
| Poisson's Ratio | 0.3 |
| **Dermis** |  |
| Density | 1.2 g/cm<sup>3</sup> |
| Young's Modulus | 0.066 MPa |
| Poisson's Ratio | 0.495 |
| Dimension (diameter x height) | 600 μm x 1000 μm [16] |
| **Epidermis** |  |
| Density | 1.3 g/cm<sup>3</sup> |
| Young's Modulus | 1 MPa |
| Poisson's Ratio | 0.495 |
| Dimension (diameter x height) | 600 μm x 100 μm [16] |

The Von Mises stress distribution was analyzed for various needle heights and pitches to evaluate the mechanical robustness of Si MNs. Heights of 500 μm and 700 μm were identified as optimal, with stress levels remaining below Si’s yield strength of 7 GPa [15]. Needles measuring 300 μm exhibited excessive stress due to poor force distribution, while 900 μm needles exceeded the yield strength, risking mechanical failure [16], as summarized in Table 2. Similarly, pitch variations of 500 μm, 1000 μm, and 1500 μm demonstrated minimal differences in stress levels, all remaining within Si’s yield strength. However, all tested configurations exceeded the minimum stress threshold of 3.18 MPa required for effective skin penetration [17]. These results highlight the importance of maintaining an adequate pitch to ensure reliable penetration while minimizing mechanical and electrical interference between adjacent needles.

**Table 2.** Analysis of different MN heights and pitches on Maximal Stress levels. The maroon highlighting indicates optimal parameters for stability and penetration effectiveness

| Parameter | Value ($\mu\text{m}$) | Maximal Stress (GPa) | Explanation |
| --- | --- | --- | --- |
| Height ($\mu\text{m}$) | 300 | 6.31 | High stress, less stable |
|  | 500 | 3.9 | Optimal stability |
|  | 700 | 3.47 | Optimal stability |
|  | 900 | 10.0 | Exceed yield strength, failure risk |
| Pitch ($\mu\text{m}$) | 500 | 3.83 | High displacement, reduces penetration |
|  | 1000 | 3.47 | Optimal |
|  | 1500 | 3.39 | Optimal |

Buckling resistance was assessed using critical load factors, where values above one indicates structural stability. A constant force of 5 N was applied along each needle’s axis [14]. While 300 μm and 500 μm needles demonstrated superior buckling performance with critical load factors of 80 and 81, the 500 μm and 700 μm needles were ultimately selected as optimal. This selection strikes a balance between mechanical stability and effective penetration depth. In contrast, 900 μm needles exhibited the lowest stability, with a critical load factor of 45, rendering them unsuitable for practical applications.

Interestingly, while needle buckling is generally considered undesirable, studies have suggested that skin buckling may enhance penetration efficiency [17][18]. This prompted an analysis of skin displacement as a function of pitch. Needles spaced at a 500 μm pitch caused greater skin displacement than those at a 1500 μm pitch, due to amplified deformation forces from closer spacing, as shown in Figure 2. However, despite the increased displacement, irregular force overlaps and stress distribution at tighter pitches made wider configurations (1000 μm and 1500 μm) more favorable for controlled and effective penetration. These findings showcase the critical role of needle spacing in achieving optimal skin-electrode interaction.

**Figure 1.**
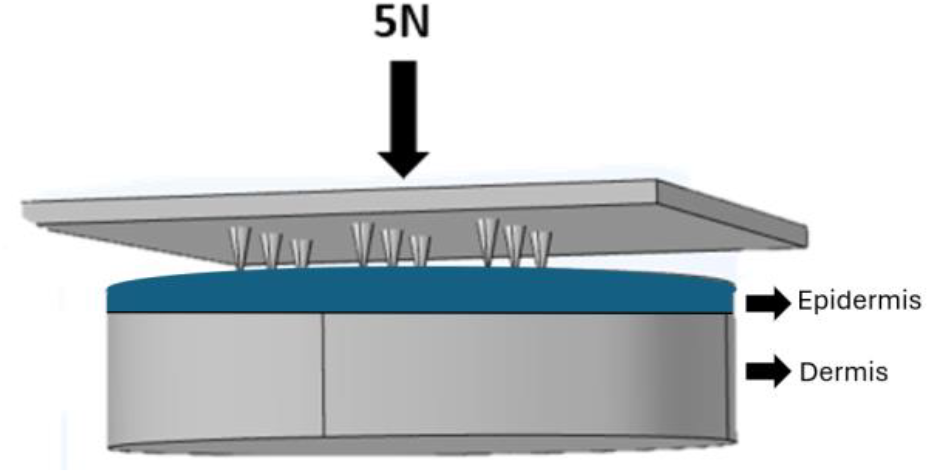
The Comsol Model

**Figure 2.**
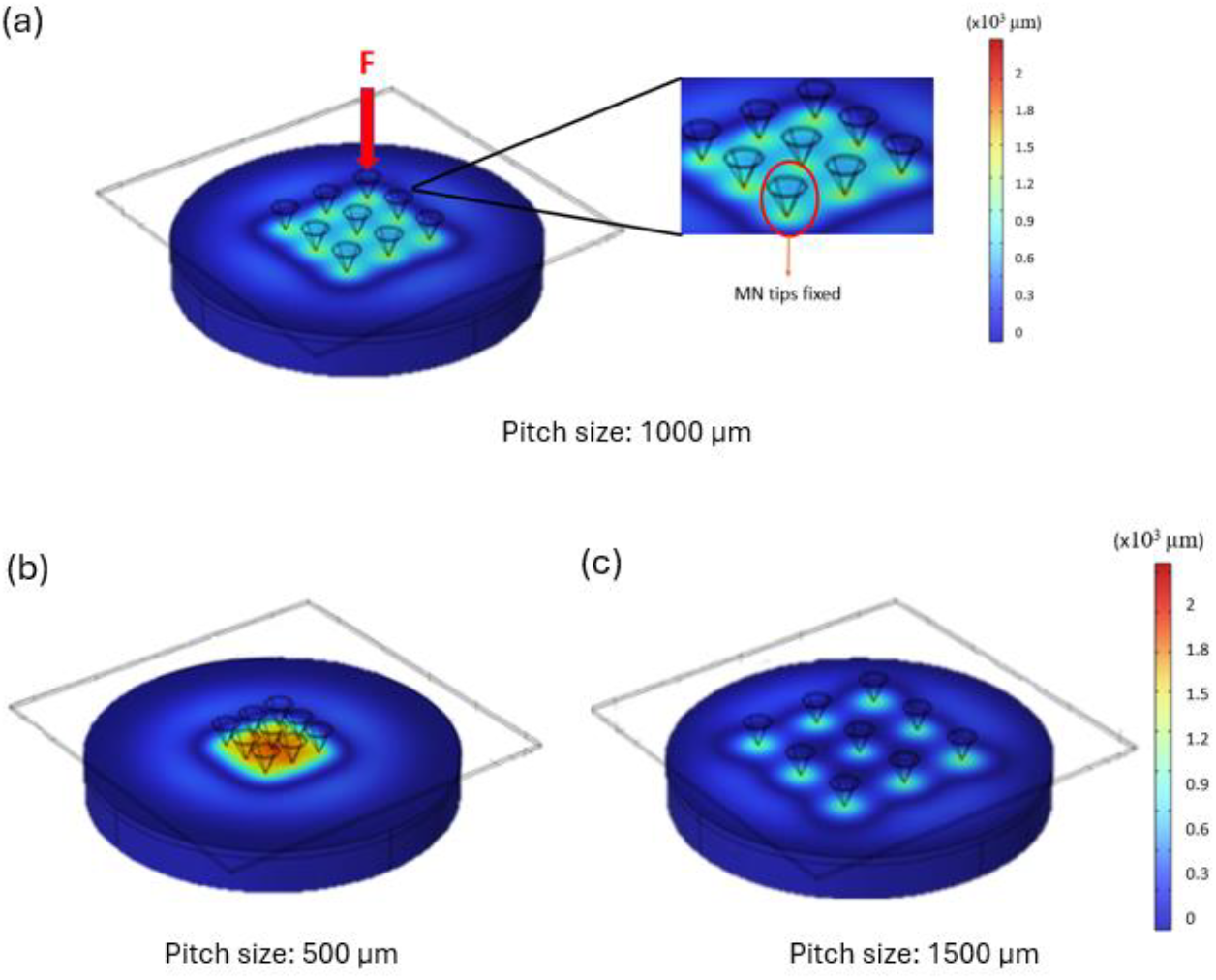
COMSOL simulations illustrating skin displacement under MN pressure for varying pitch sizes. (a) Pitch size of 1000 μm showing the force application and fixed needle tips. (b) Displacement using MNs at a 500 μm pitch, demonstrating intense local deformation. (c) Displacement using MNs at a 1500 μm pitch, showing reduced overlap and less intense deformation compared to the 500 μm pitch.

## Fabrication Process

The fabrication process, depicted in Figure 3, was streamlined based on insights from our simulations and established methods. We drew inspiration from Hasada et al. [19], who developed a multi-step process combining three dicing steps with a single wet etching step to fabricate densely packed MNs with integrated flow channels. While effective for fluidic applications, their workflow is complex. Similarly, Howels et al. [20] employed anisotropic wet etching with a nitride mask to create in-plane MNs for drug delivery, relying entirely on etching conditions to define geometry. Building on these approaches, our method simplifies the workflow by integrating a single dicing step with anisotropic wet etching. This two-step process eliminates the need for iterative dicing or lithographic patterning, providing a scalable and reproducible solution with precise control over MN geometry. If additional patterning were required, it could be achieved using masks or resist protection to define specific features during etching.

**Figure 3.**
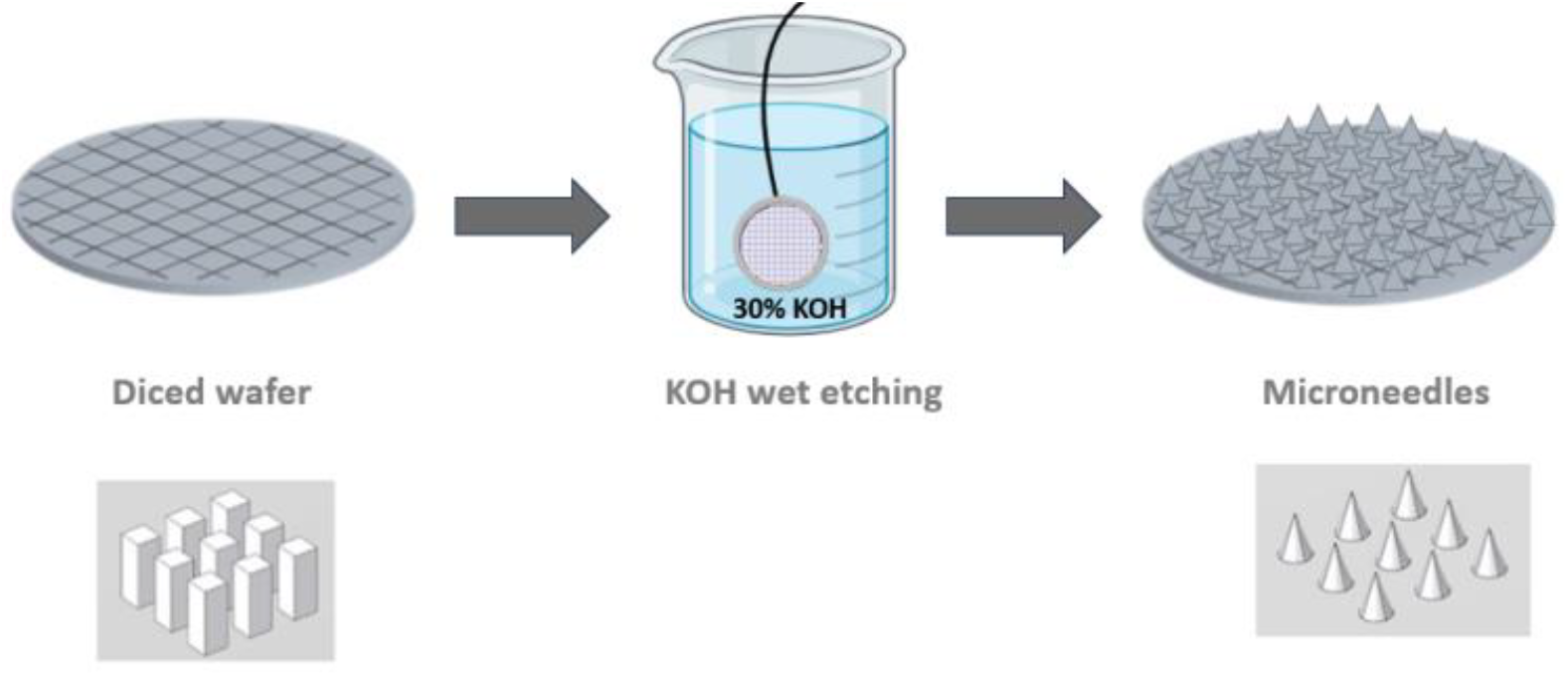
Process flow for Si MNs

Results from simulations indicated that dicing to a depth of 700 μm, followed by immersion in a 30% potassium hydroxide (KOH) solution heated to 80°C, optimizes the etch rate while minimizing the risk of undercutting [21]. Upon immersion, the wet etching process begins, with the KOH attacking the Si in the <100> plane. The nitride layer acts as a protective barrier, resulting in a characteristic anisotropic V-etch with well-defined sidewalls [20]. As the etching progresses, needles begin to form, each topped with a ‘hat’ (Figure 4). The process is considered complete when the etched silicon ‘hats’ detach from the needles.

**Figure 4.**
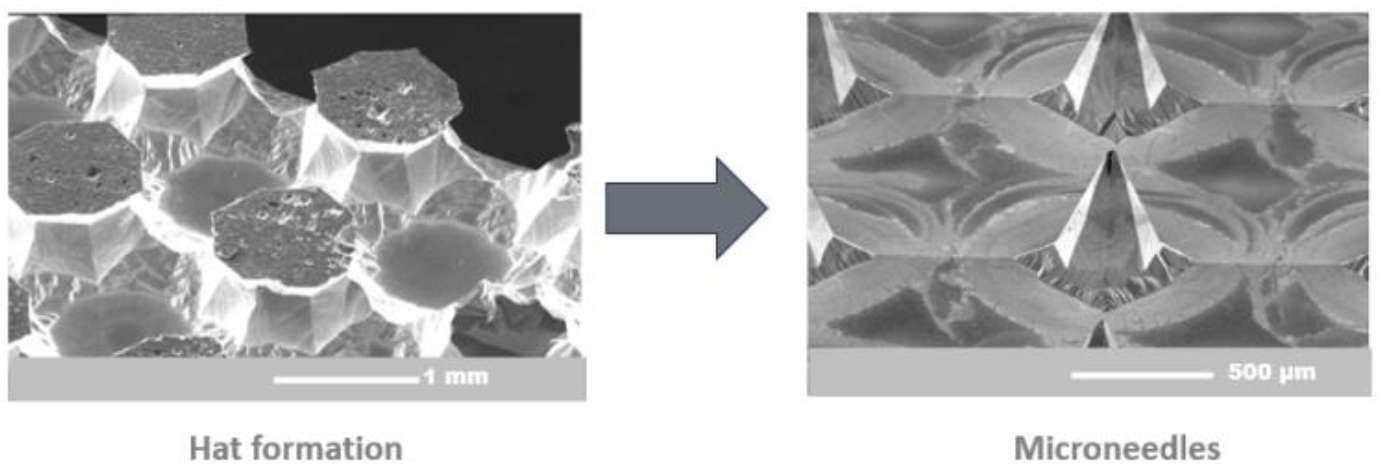
SEM images showing the process of Si MNs formation

Based on the simulation findings, and considering the practical constraints of the fabrication process, the wafer was diced with a grid distance of 1.7 mm and a cutting depth of 700 μm before undergoing wet etching. This grid distance ensures sufficient spacing to prevent mechanical interference between adjacent needles while optimizing needle density per unit area. After about nine to ten hours of etching, the ‘hats’ typically detached, revealing sharp needles as shown in Figure 4. The MNs measured a radius of 435 μm and a height of approximately 700 μm.

## EEG Validation as a Proof of Concept

To evaluate the effectiveness of the MN electrode array in recording biosignals, we tested the electrodes in an EEG recording experiment. EEG consists of a set of scalp electrodes that measure very weak biopotential signals (usually within the microvolt range) generated by the synchronized activity of large neuron populations within the brain. We conducted two experiments to record steady-state visual evoked potentials (SSVEP) [22] and auditory steady-state responses (ASSR) [23], which are neural responses that phase-lock to a periodic visual and auditory stimulus, respectively. The objective was to validate the performance of the MN electrodes in capturing SSVEPs and ASSRs. Additionally, we compared the signals acquired with our MN electrodes and with a commonly used wet electrode (Ag|AgCl with SuperVisc conductive gel). Our experiments were approved by the KU Leuven Social and Societal Ethics Committee.

The EEG measurement system consists of a miniature sensor node and a data sink node connected to a computer via USB. Each sensor node comprises a 5.5 cm × 2 cm PCB, using the ADS1299 chip [24]. Bias resistors replace the traditional right leg drive (RLD) circuit, reducing the number of required electrodes while maintaining signal quality without the need for a dedicated bias electrode. The sensor node continuously transmits its EEG data to the data sink, which synchronizes the data streams and forwards the aligned data to the computer for further analysis. Further details on the system can be found in [25].

In our study, we standardized the inter-electrode distance at 3 cm for all measurements and validations, as research has shown that distances below this threshold significantly affect the decoding quality of auditory responses [26]. By validating our system under this constrained condition, we establish its robustness in compact configurations, e.g., for wearable sensors. Given that larger inter-electrode distances typically enhance signal quality, the system’s demonstrated performance at 3 cm further substantiates its applicability to configurations with greater electrode spacing.

### (i) Impedance measurements

MN electrodes offer several practical advantages over traditional wet electrodes for EEG recordings. They do not require conductive gels or extensive skin preparation, which simplifies the setup and preparation process and reduces the risk of skin irritation. These features make MN electrodes particularly suitable for portable and wearable EEG systems, where ease of use and comfort are important considerations. Wet electrodes, with the help of conductive gels, typically achieve impedance values between 10 and 30 kΩ [27]. However, dry electrodes, such as our MN electrodes, generally exhibit higher impedance than wet electrodes.

In our experiments, each MN array consisted of 16 needles, corresponding to a total electrode area of 7×7 mm. Both the MN electrodes and the commercial Ag|AgCl wet electrodes (size: 9 mm diameter) were positioned on the human scalp with a 3cm spacing within their respective pairs. To evaluate and compare the impedance of different electrode types, we conducted impedance measurements across a wide frequency range (20 Hz to 100 kHz) using the Wayne Kerr 6500B impedance analyzer. While the EEG frequency range typically spans from 0.5 Hz to 100 Hz [27], the measurements were limited to 20 Hz and above due to the equipment’s specifications. The electrodes were attached to the scalp using surgical tape, which ensured proper placement throughout the measurement process. The results showed that the bare Pt MN electrodes had an impedance of approximately 95.5 kΩ at 20 Hz, higher than the Ag|AgCl electrodes, which maintained lower impedance throughout the frequency range as shown in Figure 5. Despite this higher impedance, the MN array still provided reliable signal detection (see the next test results), even with a close spacing of 3 cm between the electrodes placed on the skin.

**Figure 5.**
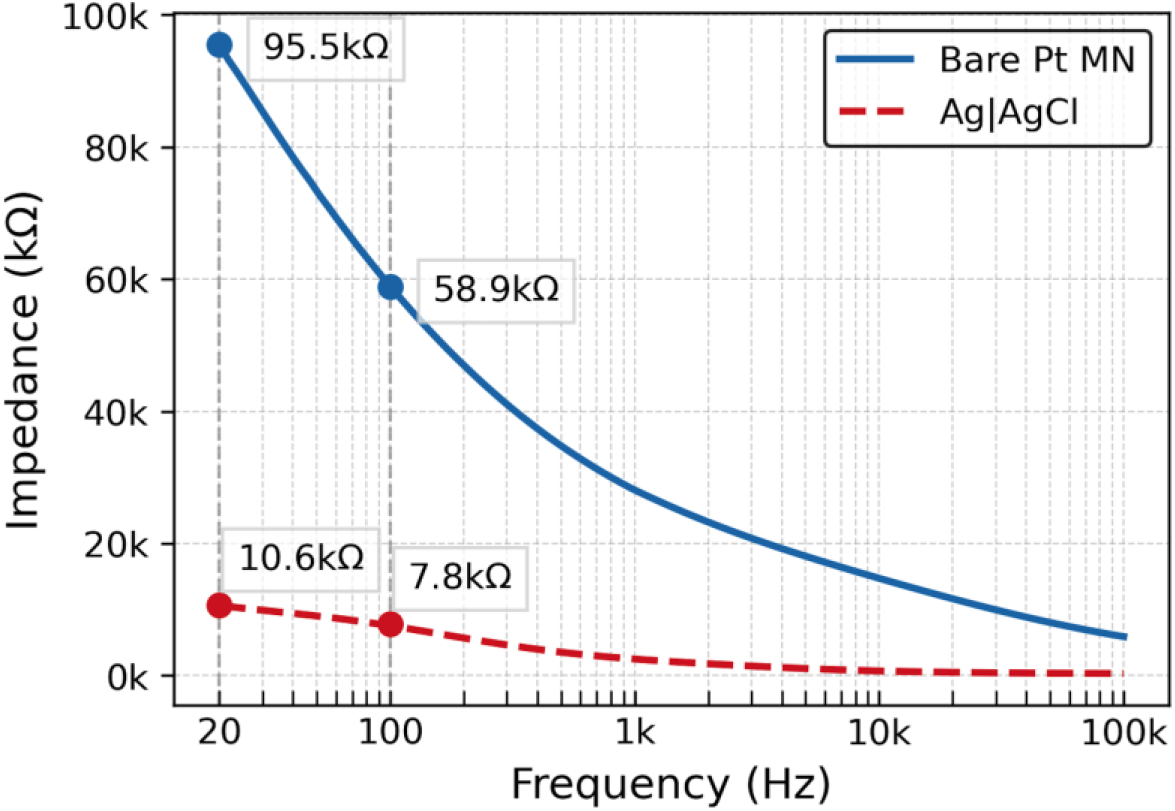
Comparison of impedance between bare Pt MN electrodes and Ag|AgCl wet electrodes on human skin

### (ii) SSVEP validation

A steady-state visual evoked potential (SSVEP) is a neural response generated when the brain is exposed to a repetitive visual stimulus, such as a flickering screen [22]. This response occurs at the same frequency as the stimulus and can be detected in EEG signals recorded over the visual cortex. SSVEPs are widely used in brain-computer interface (BCI) applications due to their well-defined spectral characteristics and high signal stability. In the SSVEP experiment, we employed a 2-electrode montage. Based on the extended 10-20 System for EEG electrode placement, the active signal measurement electrode was positioned between O1 and Oz, over the visual cortex, while the reference electrode was placed between POz and Oz [28]. These positions are shown in Figure 6, with “EEG Signal” as the active electrode and “REF” as the reference electrode.

**Figure 6.**
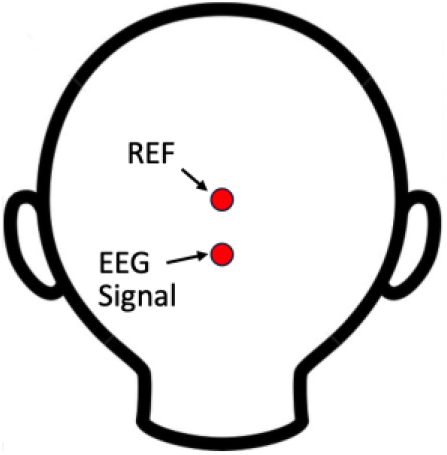
Positions for the 2-electrode SSVEP montage

Continuous EEG data was recorded for one minute, while the subjects were watching a flickering screen switching between black and white at a frequency of 12.5 Hz. As a result, the SSVEP will be visible in the EEG spectrum as a narrow peak at 12.5 Hz (and its harmonics). Figure 7 presents the EEG power spectrum obtained from a single measurement using the MN electrodes. The spectrum clearly demonstrates a distinct peak at 12.5 Hz, corresponding to the flickering stimulus frequency. This result highlights the ability of the MN electrodes to reliably capture SSVEP signals. After computing a periodogram (i.e., squared magnitude of the discrete Fourier transform) over 1 minute of EEG data, the signal-to-noise ratio (SNR) was calculated as the ratio of the 12.5 Hz target peak to the average noise level between 5 Hz and 20 Hz.

**Figure 7.**
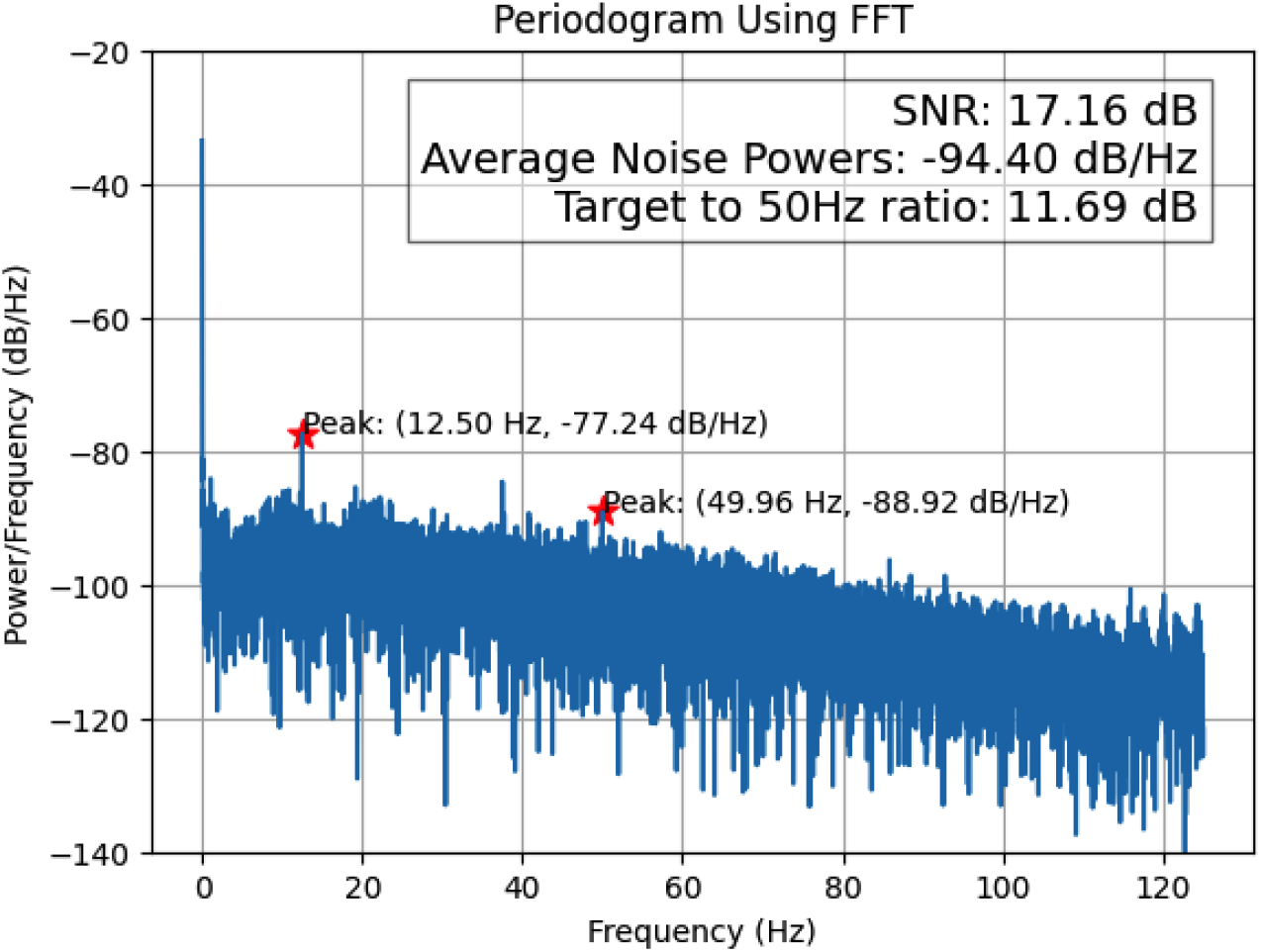
EEG power spectrum from a single SSVEP measurement using the MN electrodes

The results summarized in the box plot in Figure 8 present the SNR values from 15 measurements obtained from a single subject using both the bare Pt MN electrodes and the Ag|AgCl wet electrodes. These measurements were conducted on five separate days, with three recordings per day. The box plot was generated from these measurements, with the box representing the interquartile range (IQR) and the red line indicating the median SNR value. Whiskers extend to the extreme data points within 1.5 × IQR, and outliers are marked as blue dots. The wet electrodes achieved a higher median SNR (21.84 dB) compared to the barePt MN electrodes (17.39 dB). Despite this difference, the bare Pt MN electrodes still achieved sufficient SNR for detecting the 12.5 Hz target signal, demonstrating their capability for reliable signal acquisition. These results suggest that while wet electrodes may be optimal for scenarios requiring maximum signal clarity, the bare Pt MN electrodes provide a practical alternative in applications prioritizing portability, ease of use, or long-term monitoring.

**Figure 8.**
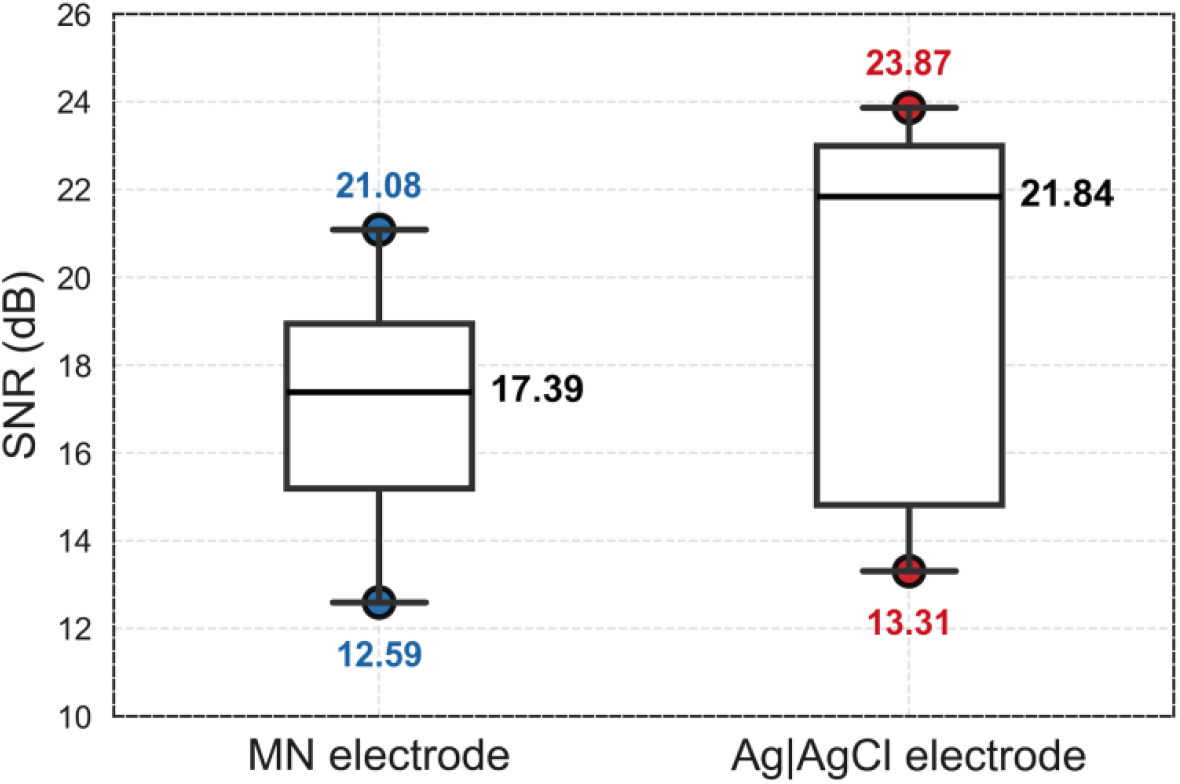
Comparison of SNR for bare Pt MN and Ag|AgCl electrodes during SSVEP experiment

### (iii) ASSR validation

To evaluate the system’s effectiveness for a different stimulus-response paradigm, auditory steady-state response (ASSR) experiments were conducted on three participants (1 male, 28 years old; 2 females, 28 and 30 years old). ASSRs are periodic neural responses elicited by amplitude-modulated auditory stimuli, reflecting brain activity synchronized to the stimulus modulation frequency [23]. These responses are widely used in auditory neuroscience and clinical applications to assess auditory processing and brainstem function. Participants were presented with binaural 40 Hz ASSR stimuli (1 kHz carrier tone, 100% amplitude modulation) via insert headphones (calibrated at a comfortable listening level). EEG signals were recorded using our custom-designed EEG system, with electrodes placed behind the right ear and referenced to the right earlobe. Both barePt MN electrodes and Ag|AgCl wet electrodes were positioned at the same location to ensure consistency. Each participant completed three 2.5-minute trials.

Figure 9 presents the distribution of SNR values for 40 Hz ASSR measurements across three subjects, comparing barePt MN electrodes and Ag|AgCl wet electrodes. Each subject completed three trials, resulting in a total of nine measurements per electrode type. The box plot represents the variability in the recorded SNR values, with the central line indicating the median SNR and whiskers extending to data points within 1.5 times the interquartile range. Outliers are displayed as individual markers. The Ag|AgCl wet electrodes exhibited a higher median SNR than the barePt MN electrodes, with a slightly broader distribution across trials. While the wet electrodes provided improved SNR, the barePt MN electrodes remained capable of reliable 40 Hz ASSR detection.

**Figure 9.**
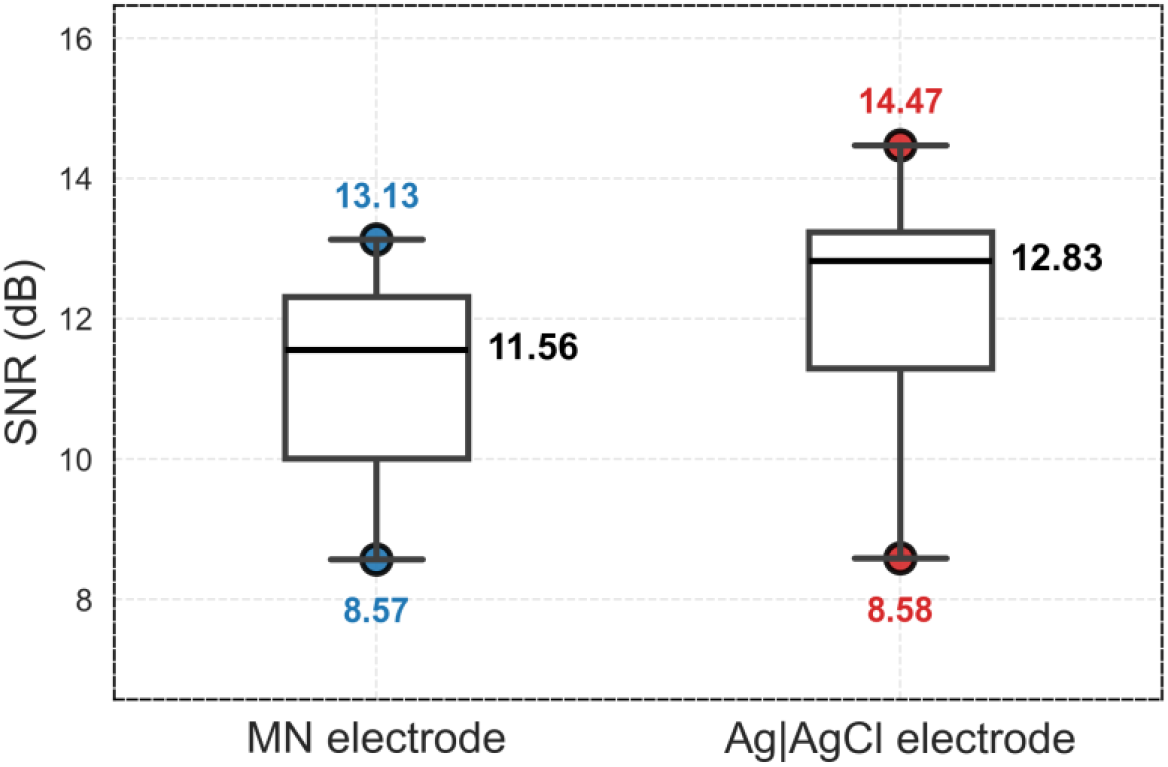
Comparison of SNR for barePt MN and Ag|AgCl electrodes during ASSR experiment

Overall, these experiments suggest that while wet electrodes remain preferable for applications requiring high signal fidelity, the barePt MN electrodes offer a viable alternative for scenarios emphasizing portability, ease of use, and long-term monitoring.

## Perspectives

In addition to the barePt MN electrodes, we developed nanostructured platinum (nanoPt) MN electrodes through an electrodeposition process, which enhances the electrochemically active surface area by modifying the platinum layer at the microstructure level. This increased surface roughness is expected to lower impedance and improve EEG signal acquisition.

To preliminarily assess their performance, we conducted impedance measurements on a skin-mimicking agarose gel. The results showed that nanoPt MN electrodes exhibited significantly lower impedance compared to barePt MN electrodes, with values of 285.4 Ω at 20 Hz, representing a reduction of approximately 41% from the 486.3 Ω measured for the barePt MN electrodes. This suggests enhanced electrical conductivity and potential benefits for neurophysiological recordings compared to bare platinum MN electrodes. While agarose gel does not fully replicate human skin, these initial findings indicate that nanostructuration could improve electrode performance for EEG applications. Further validation on human subjects is required to confirm these findings and assess long-term usability. As ethical approval has not yet been obtained, in vivo testing will be pursued once the necessary approvals are secured. These tests will evaluate the efficacy of nanoPt MN electrodes compared to commercial alternatives. If confirmed, the reduced impedance and improved signal quality could make nanoPt MNs a promising solution for portable and wearable EEG systems, offering a reliable and minimally invasive interface for enhanced neurophysiological monitoring.

## Conclusion

This study presents a streamlined, two-step fabrication approach for Si MN electrode arrays designed for bio-signal acquisition, with a focus on EEG applications. Finite element modeling using COMSOL Multiphysics guided the optimization of the MN array, ensuring mechanical robustness and minimizing skin-electrode impedance. Structural simulations identified an optimal needle length range of 500–700 μm, balancing mechanical strength and effective skin penetration while preventing buckling. Needle spacing was also optimized to minimize inter-needle interference and enhance signal acquisition efficiency.

Experimental validation demonstrated that the fabricated platinum-coated Si MN electrodes achieved stable EEG recordings comparable to conventional wet electrodes while offering the practical benefits of dry electrodes. By eliminating the need for conductive gels and skin preparation, MN electrodes reduce setup time and enhance user convenience, making them particularly suitable for wearable and long-term monitoring applications. The proposed fabrication method is cleanroom-compatible, scalable, and cost-effective compared to more complex microfabrication techniques. These results establish Si MN electrodes as a viable solution for dry-electrode EEG applications. Future work will focus on acquiring more data for ‘on-skin’ validation and benchmarking against commercial alternatives. In addition, these tests will be conducted using nanostructured electrodes created with one-step, cleanroom-compatible methods and biocompatible materials, which will further advance bio-signal monitoring technologies.

## Acknowledgements

This project has received funding from the European Research Council (ERC) under the European Union’s Horizon 2020 research and innovation programme (grant agreement No 802895 and 101138304). Views and opinions expressed are, however, those of the author(s) only and do not necessarily reflect those of the European Union or ERC. Neither the European Union nor the granting authority can be held responsible for them. The authors express gratitude to the Nanostacks project for financial support under Grant Agreement No. 951949 as well as to KU Leuven (and the Industrieel Onderzoeksfonds (C3 project-3E230062).

## Notes

### Competing Interest Statement

The authors have declared no competing interest.

